# How many digits in a mean are worth reporting?

**DOI:** 10.1101/003558

**Authors:** R. S. Clymo

**Affiliations:** School of Biological and Chemical Sciences, Queen Mary University of London, London E1 4NS, UK

## Abstract

Most bioscientists need to report mean values, yet many have little idea of how many digits are significant, and at what point further digits are mere random junk. Thus a recent report that the mean of 17 values was 3.863 with a standard error of the mean (SEM) of 2.162 revealed only that none of the seven authors understood the limitations of their work. The simple rule derived here by experiment for restricting a mean value to its significant digits (sig-digs) is this: the last sig-dig in the mean value is at the same decimal decade as the first sig-dig (the first non-zero) in the SEM. An extended rule for the mean, and a different rule for the SEM itself are also derived. For the example above the reported values should be a mean of 4 with SEM 2.2. Routine application of these simple rules will often show that a result is not as compelling as one had hoped.

## Introduction

During the last five years, as referee (reviewer) or editor of 53 biosciences articles, I found that 29 (55 %) reported too many, even ridiculously too many, digits in mean values or SEMs or both. For example, a report of 3.863 ± 2.162 for a sample of 17 for which the sig-digs are really 4 ± 2.2. A scan of 50 articles in a variety of bioscience journals showed that 32 % made this mistake once or many times. It is not the total number of digits, or where the decimal point falls that matters: the critical feature is the relation between the mean and its SEM. Thus the frequency of a transition of a trapped and laser-cooled, lone ion of ^88^Sr^+^ was reported^1^, correctly according to the extended rule in Table 3, to 16 significant digits as 444 779 044 095 484.6 Hz, with a SEM of 1.5 Hz. Equally correct though less clear would have been 444 779 044 .095 4846 MHz and SEM 0.000 00015 MHz.

The problem seems to be that there is no published logical analysis of where to stop. Here I derive simple rules by experiment which allow one to restrict a mean and SEM to their sig-digs: that is to those digits that do mean something and are not just random junk.

## Experiments and their results

From a population with mean 39.615 and SEM 1.33, 8000 instances were drawn at random by a computer program (8000 is an arbitrary large number). Table 1 below shows the frequency of the ten digits in successive decades. The digit ‘3’ in the 10’s decade is clearly meaningful, and so is ‘9’ in the 1’s, but in the 0.1’s decade, the target digit ‘6’, though the most frequent in its decade, is barely better than random: a mean of ‘39’ is worth reporting, but ‘39.6’ is overdoing things.

**Table 1.** Frequency of digits in 8000 values drawn at random from a ‘normal’ (Gaussian) population with mean 39.615, SEM 1.33. The target digit in each decade is <u>underlined</u>.

| Decade | Digit |  |  |  |  |  |  |  |  |  |
| --- | --- | --- | --- | --- | --- | --- | --- | --- | --- | --- |
|  | 0 | 1 | 2 | 3 | 4 | 5 | 6 | 7 | 8 | 9 |
| 10's | ... | ... | ... | <u>4846</u> | 3154 | ... | ... | ... | ... | ... |
| 1's | 1963 | 890 | 261 | 36 | 5 | 33 | 190 | 677 | 1621 | <u>2324</u> |
| 0.1's | 827 | 798 | 787 | 775 | 769 | 777 | <u>859</u> | 810 | 840 | 758 |

Table 2 below derives from the same mean but a SEM 100 times smaller at 0.0133. This supports a mean of 39.61, but in the next ‘0.00x’ decade the target digit, ‘5’, is not even the most frequent in its decade. **A simple rule is, thus, to stop the mean at the same decade as that of the *first* significant (non-zero) digit in the SEM**. Note that the rule uses the SEM to show where to stop: it makes no use whatever of the position of the decimal point.

**Table 2.** As in Table 1, but with SEM 0.0133, only 1/100 of that in Table 1. In the last row the target digit (‘5’, in 39.615) is not even the most frequent digit.

| Decade | Digit |  |  |  |  |  |  |  |  |  |
| --- | --- | --- | --- | --- | --- | --- | --- | --- | --- | --- |
|  | 0 | 1 | 2 | 3 | 4 | 5 | 6 | 7 | 8 | 9 |
| 10's | ... | ... | ... | <u>8000</u> | ... | ... | ... | ... | ... | ... |
| 1's | ... | ... | ... | ... | ... | ... | ... | ... | ... | <u>8000</u> |
| 0.1's | ... | ... | ... | ... | ... | 924 | <u>7076</u> | ... | ... | ... |
| 0.01's | 1738 | <u>2331</u> | 1825 | 894 | 240 | 43 | 5 | 35 | 164 | 725 |
| 0.0001's | 759 | 823 | 787 | 806 | 800 | <u>768</u> | 797 | 827 | 813 | 820 |

In Table 1 the counts in the ‘0.1’s decade are near random, but if we were to decrease the SEM gradually the totals for each digit in a decade would become more and more unequal as peaks emerged and grew from the slowly sinking hummocky plain and, consequently, indicated that we would soon be able to justify another sig-dig. In a report, the number of sig-digs must be integer, but to understand the trends we need a sig-dig index, *D*_M_, that is at least semi-continuous. Such an index is derived in the Appendix.

The points in Figure 1 below show how *D*_M_ depends experimentally on *C*, the quotient of mean / SEM in experiments similar to those outlined in Tables 1 and 2. The fitted line is *D*_M_ = log_10_ (*C*).

**Figure 1.**
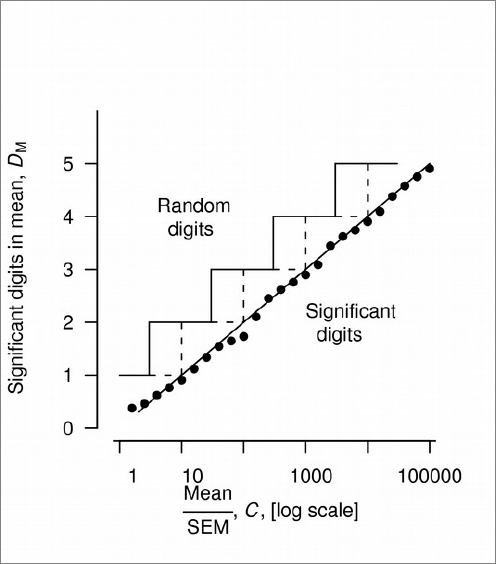
Experimental dependence (filled circles) of sig-digs in a mean on the *C* = mean / SEM quotient. The points are close to the calculated D_M_ = log_10_ (C) line. The area below and to the right of the sloping line is the domain of significant digits in the mean; above and to the left the digits are random junk. The broken line staircase shows the simple rule for integer (1, 2, 3 and so on) sig-digs. The unbroken line staircase shows a better but slightly more complex rule that gives a more uniform distance between the staircase and the continuous line. Both simple and extended rules are shown in Table 3.

If we take the ceiling of these values — equivalent to truncating and adding 1 — to get an integer value we get the stepped broken line in Figure 1. This translates into the simple rule already given. But at some points, at the back of the steps, this gives values that are only just sufficient while at the front of the step the values are well into the meaningless random zone and almost a digit too many. A more complex extension to the simple rule (Table 3) shifts the steps about half a decade left (log_10_ ≈ (3) 0.5) and spreads the overshoot into the meaningless region more evenly: the overshoot is 0.5 to 1.5 random digits. The unbroken line (staircase) in Figure 1 shows the extended rule.

**Table 3.**
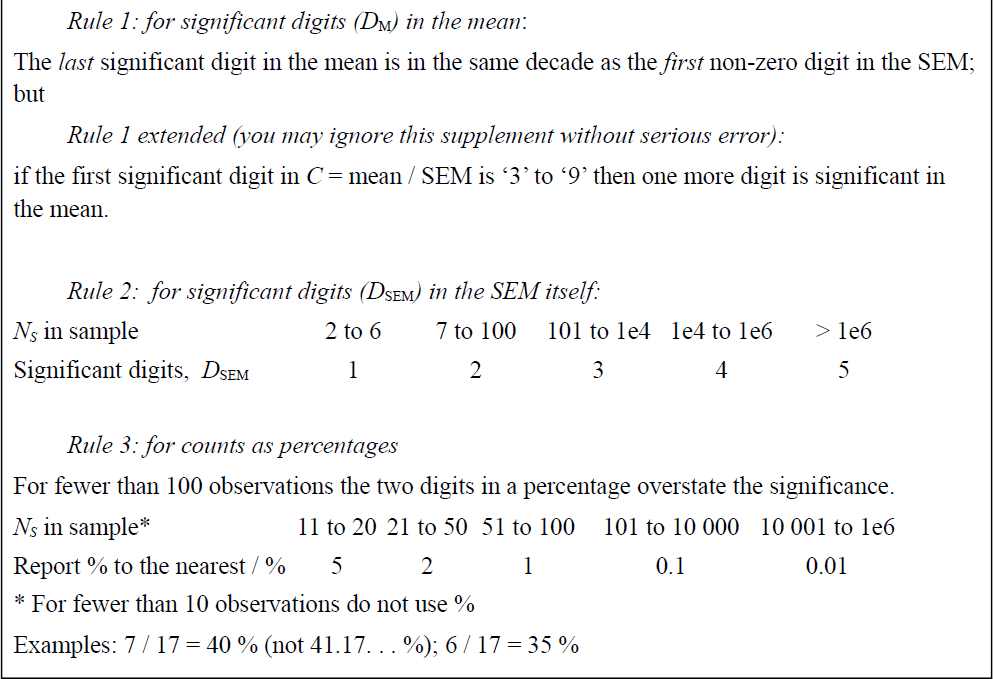
Rules determining the number of significant digits to report

|  |  |  |  |  |  |
| --- | --- | --- | --- | --- | --- |
| $N_S$ in sample | 2 to 6 | 7 to 100 | 101 to 1e4 | 1e4 to 1e6 | > 1e6 |
| Significant digits, $D_{\text{SEM}}$ | 1 | 2 | 3 | 4 | 5 |
| $N_S$ in sample* | 11 to 20 | 21 to 50 | 51 to 100 | 101 to 10 000 | 10 001 to 1e6 |
| Report % to the nearest / % | 5 | 2 | 1 | 0.1 | 0.01 |

\* For fewer than 10 observations do not use %
Examples: $7 / 17 = 40\%$ (not 41.17... %); $6 / 17 = 35\%$

Rule 2 for *D*_SEM_ is simpler but its origin is more complicated. Figure 2 below shows, for a fixed mean and standard deviation (SD), how *D*_SEM_ depends, in experiments similar to those in Tables 1 and 2, on the number of items, *N*_s_, in the calculation of a SEM. Points for two such experiments, with the same mean and different SDs are shown. Over a range of 100 in *N*_s_ the value of *D*_SEM_ rises with a slope of 1 on the log-linear scales shown: *D*_SEM_= log_10_ (*N*_s_) + *c.* But eventually it falls over a cliff from a first significant digit of ‘1’ to ‘9’ a decade further down (for example from 1.001 to 0.999). The overall slope of this saw-toothed progression (0.5) is half that of the teeth themselves reflecting the fact that the SEM depends on √*N*_s_.

**Figure 2.**
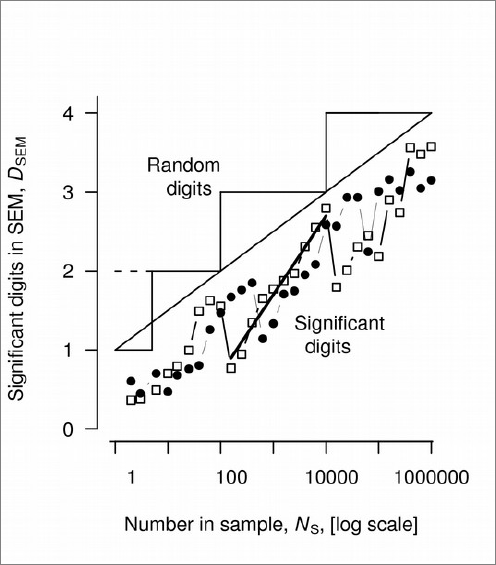
For a fixed standard deviation (SD), the experimental dependence (filled circles) of sig-digs in the SEM, on the number, *N*_S_, of items in the sample on which the SEM is based. The points are close to a series of segments D_SEM_ = log_10_ (N_s_) + c. The saw-tooth jumps occur where the first significant digit of the SEM, which depends on N_s_, passes from 1 to 9 (e.g. from 1.001 to 0.999). The unfilled squares show a similar pattern for a SD half that for the filled circles. The line through the points for one sawtooth has a slope of 1.0. The longer sloping line y = log_10_ (N_s_) / 2 + 1, with half the slope of the saw-tooth lines, summarises the upper bound of saw-tooth lines and sets the boundary between significant and random digits. The staircase ending in a broken line, with a step every 100-fold increase in Ns shows the simple rule for significant digits in a SEM. The staircase with continuous lines and a short step at the bottom shows the more complex rule listed in Table 3, taking account of behaviour at small N_s_.

The exact position of the saw-tooth depends on the numerical value of the SEM, and to accommodate this the bounding line *D*_SEM_ = log_10_ (*N*_s_) / 2 + 1 is shown. The steps show Rule 2. The offset for *N*_s_ ≤ 6 accommodates the fact that at small *N*_s_ the bounding line curves downwards, though this is not shown in detail in Figure 2.

The full rules for sig-digs in a mean and in a SEM are summarised below in boxed Table 3, which includes a rule for percentages (which have additional problems).

A minor problem remains. Suppose that a computer-calculated mean is 12345.67 with calculated SEM 5678.90. The results are based on 25 values, so the SEM justifies 2 sig-digs. These would be ‘57’, but how can we show where the decimal point is? One way would be to change units, if there are any. A more general solution though is to use italic ‘0’ as a *packing digit*. The example would then be reported as 12000 with SEM 5700.

## Discussion

Does it matter if random junk is reported? For those who understand these matters, no. They can adjust the values themselves. For an author’s reputation, yes, it does matter. Gross over-reporting of values is one of the clearest warning signs to readers that the author does not understand what he or she is doing.

The big general purpose journals were not interested in this article, yet (I suggest) those who need it most are the least likely to come across it in a specialist education or statistics journal. This version was therefore archived as: Clymo RS (2012) How many digits in a mean are worth reporting? http://arXiv.org/abs/1301.1034. But that location is unlikely to be scanned by bioscientists. ‘Duplicate publication’ is strongly deprecated in formal journals, for obvious reasons, but is allowed in preprint archives. So this is the second source for this article.

If you found it useful, please recommend it to colleagues and to editors.

## Acknowledgements

I thank all those whose reporting of junk digits irritated me enough to write this article.

## Appendix: derivation of the sig-dig index, *D*_*M*_

To calculate the sig-dig index, *D*_*M*_, first, set a threshold, *T*. Let *n* be the total number of counts. The expected number, *E*, in each of the 10 classes (the digits 0 to 9, Tables 1 and 2) if the counts are uniformly distributed is *n* / 10. Let |*X*| be an absolute difference from *E*, then the smallest χ^2^ results if half the ten classes (of the digits 0 to 9) are *E* + |*X*| and half are *E* − |*X*|. Substituting these values in the formula for calculating χ^2^ in a contingency table gives |*X*| = √(*n* χ^2^) / 10. This value of |*X*| defines the threshold, *T*. For *P* = 0.5 and 9 d.f., χ^2^ = 8.34. Any value with |*X*|, the difference from *n* / 10, < 0.289 √*n* is more likely to be random than significant.

The maximum count above the expected value is *n − E*. Start at the decade in the mean value with the first non-zero digit and work rightwards through the decades (downwards in Tables 1 and 2) stopping when the counts for the decade as a whole are more likely than not to be random in the χ^2^ test at *P* = 0.5, 9 d.f. Then calculate the index as the sum, for all the decades and digit classes where the count, *x*, exceeds the threshold, *T*, of *D*_M_ = (*x* − *E*) / (*n* − *E*).

As illustrations: if all counts in a decade are for a single digit, 1.0 is added to *D*_M_. If there are equal counts for two digits in the decade and zero for the other eight, then 0.89 is added.

